# Consideration of genetic variation and evolutionary history in future conservation of Indian one-horned rhinoceros (*Rhinoceros unicornis*)

**DOI:** 10.1101/2022.01.11.475781

**Authors:** Tista Ghosh, Shrewshree Kumar, Kirtika Sharma, Parikshit Kakati, Amit Sharma, Samrat Mondol

**Author notes:** Corresponding authors: Samrat Mondol, Ph.D., Animal Ecology and Conservation Biology Department, Wildlife Institute of India, Chandrabani, Dehradun, Uttarakhand 248001. Email-, Tista Ghosh, M.Sc., Animal Ecology and Conservation Biology Department, Wildlife Institute of India, Chandrabani, Dehradun, Uttarakhand 248001.

## Abstract

The extant members of the Eurasian rhino species have experienced severe population and range declines through a combination of natural and anthropogenic factors since Pleistocene. The one-horned rhino is the only Asian species recovered from such strong population decline but most of their fragmented populations in India and Nepal are reaching carrying capacity. Implementation of any future reintroduction-based conservation efforts would greatly benefit from currently unavailable detailed genetic assessments and evolutionary history of these populations. We sequenced wild one-horned rhino mitogenome from all the extant populations (n=16 individuals) for the first time, identified the polymorphic sites and assessed genetic variation (2531bp mtDNA, n=111 individuals) across India. Results showed 30 unique rhino haplotypes distributed as three distinct genetic clades (F_st_ value 0.68-1) corresponding to the states of Assam (n=28 haplotypes), West Bengal and Uttar Pradesh (both monomorphic). Phylogenetic analyses suggest earlier coalescence of Assam (∼0.5 Mya) followed by parallel divergence of West Bengal and Uttar Pradesh/Nepal (∼0.06-0.05Mya), supported by the paleobiogeographic history of the Indian subcontinent. Combined together, we propose recognising three ‘Evolutionary Significant Units (ESUs)’ of Indian rhino. As recent assessments suggest further genetic isolations of Indian rhinos at local scales, future management efforts should focus on identifying genetically variable founder animals and consider periodic supplementation events while planning future rhino reintroduction programs in India. Such well-informed, multidisciplinary approach is the only way to ensure evolutionary, ecological and demographic stability of the species across its range.

## 1. Introduction

The members of Rhinocerotidae family were once one of the most diverse and widely distributed terrestrial herbivores with complex evolutionary history (Liu et al., 2021). By late Pleistocene, this family was reduced to only nine species (from more than 100 species) spread across Eurasia (seven species) and Africa (two species) (Wan and Zhang, 2017; Liu et al., 2021). Subsequently, early Holocene global warming (after Last Glacial Maxima) triggered their extinction in western Eurasia and southward movement of eastern Eurasian rhinos, leading to their distribution across Southeast Asia (Li et al., 2015; Wan and Zhang, 2017). Further, the range of all Eurasian rhino species (Javan, Sumatran and One-horned rhino) were affected by a combination of natural and anthropogenic factors during Pleistocene-Holocene transition period (0.015-0.009 million years ago (Mya)) (Alroy, 2011; Li et al., 2015; Lal, 2016; Mays et al., 2018; Liu et al., 2021), followed by recent events of exploitation of natural resources (during colonial era), industrialisation and poaching (since 17^th^ century) (Menon 1996; Fernando et al., 2006; Das et al., 2015; Steiner et al., 2017). Population size of the most widely distributed Javan rhinos (during Holocene) (Antoine, 2012) were greatly reduced during human population expansion since 10,000 years ago (Li et al., 2015), whereas the Sumatran rhino populations became fragmented and isolated (since Holocene) due to submerged Sundaland corridors (late Pleistocene) (Mays et al., 2018). The one-horned rhinos faced climate-change driven habitat shrinkage in late Pleistocene (Patnaik, 2016). Currently the Javan and Sumatran rhinos are categorized as Critically Endangered (∼60 Javan rhino - Margaryan et al., 2020 and <100 Sumatran rhinos- Steiner et al., 2017) and one-horned rhino as Vulnerable by IUCN (∼3700 individual, Talukdar, 2021). Recovery of these species in their natural habitats requires deeper understanding of demography, ecology and genetics for appropriate conservation measures.

The one-horned rhino, being the only Asian species recovered from severe population decline in the past are critical for the evolutionary potential of this group. With a current population size of ∼3700 individuals (increased from few hundred individuals in 1990s), it retains ∼96% of the Asian rhino population (Steiner et al., 2017; Margaryan et al., 2020; Talukdar, 2021). As majority of the current one-horned rhino bearing areas in India and Nepal are reaching to their carrying capacities (Subedi et al., 2017; Jhala et al., 2021), future conservation efforts are directing towards reintroduction-based programmes. Detailed genetic assessment of the existing rhino populations is critical in this regard since strong historical demographic declines has led to loss of genetic variation in all rhino species (Black rhino - Moodley et al., 2017, White rhino - Moodley et al., 2018, Sumatran rhino-Mays et al., 2018 , Javan rhino - Margaryan et al., 2020). For example, Liu et al., (2021) suggested low population size and reduced genetic diversity across Rhinocerotidae family for an extended period of time. Similarly, mitogenome-based phylogeography reported low variations in both Sumatran (Steiner et al., 2017) and Javan (Margaryan et al., 2020) rhinos, but no such data is available for one-horned rhinos.

In this paper, we investigated the phylogeography and evolutionary history of one-horned rhinos in India (henceforth Indian rhino) as it harbours 83% (Kakati et al., 2021) of the global population of this species. We sequenced all the polymorphic sites in the Indian rhino mitogenome in 111 wild individuals surveyed across seven extant populations covering the states of Assam, West Bengal and Uttar Pradesh. Further, we identified the Evolutionary Significant Units (ESUs) in Indian rhinos and suggested appropriate conservation measures to secure the evolutionary potential of this species. We believe that the results will provide the most exhaustive genetic information for Indian rhinos that would be useful in future reintroduction and population management efforts.

## 2. Materials and Methods

### 2.1 Permission and ethical considerations

Data generated in this study is part of a collaborative programme titled “Implementing Rhino DNA Indexing System to counter rhino poaching threat and aid population management in India” (henceforth RhoDIS-India). Biological sampling from all the three rhino bearing states was permitted by Ministry of Environment, Forests and Climate Change (MoEF&CC), Government of India (Letter No. 4-22/2015/WL). Permission for dung sampling was provided by state forest departments of Assam (Letter No. A/GWL/RhoDIS/2017/913, 3653/WL/2W-525/2018, WL/FE.15/22), West Bengal (Letter No. 3967/WI/2W-525/2018) and Uttar Pradesh (Letter No. 1978/23-2-12 (G)). We have also received one tissue sample from Valmiki National Park, Bihar forest department assumed to be representing the wild rhinos of Nepal (Letter/no.-1296 dated 16.10.2020). No ethical permissions were required for tissues as they were collected from naturally dead rhinos as well as for dung samples.

### 2.2 Sample collection

In this study we have used 160 samples (72 tissues and 88 dung) covering all the seven rhino bearing parks of India (Fig. 1). The tissue samples of naturally dead rhino were provided by respective forest departments as part of RhoDIS-India protocol (2017-2021). Further dung collection was done to ensure spatial coverage for areas with no representative tissue samples. Rhino dung sampling can be challenging in the wild due to their use of communal latrine system (middens) (Laurie, 1982; Johnsingh and Manjrekar, 2016). In this study, sampling was conducted by intensive foot and vehicle surveys from already known midden sites across six rhino bearing parks (except Kaziranga NP). During sampling, only the fresh bolus from top of the midden was selected and swabbed twice with separate PBS-soaked sterile cotton swabs (Himedia, Mumbai, India). All samples were geo-tagged and transferred to laboratory in -20°C freezer till downstream processing. Park-wise sample details are provided in Table A1.

**Figure 1:**
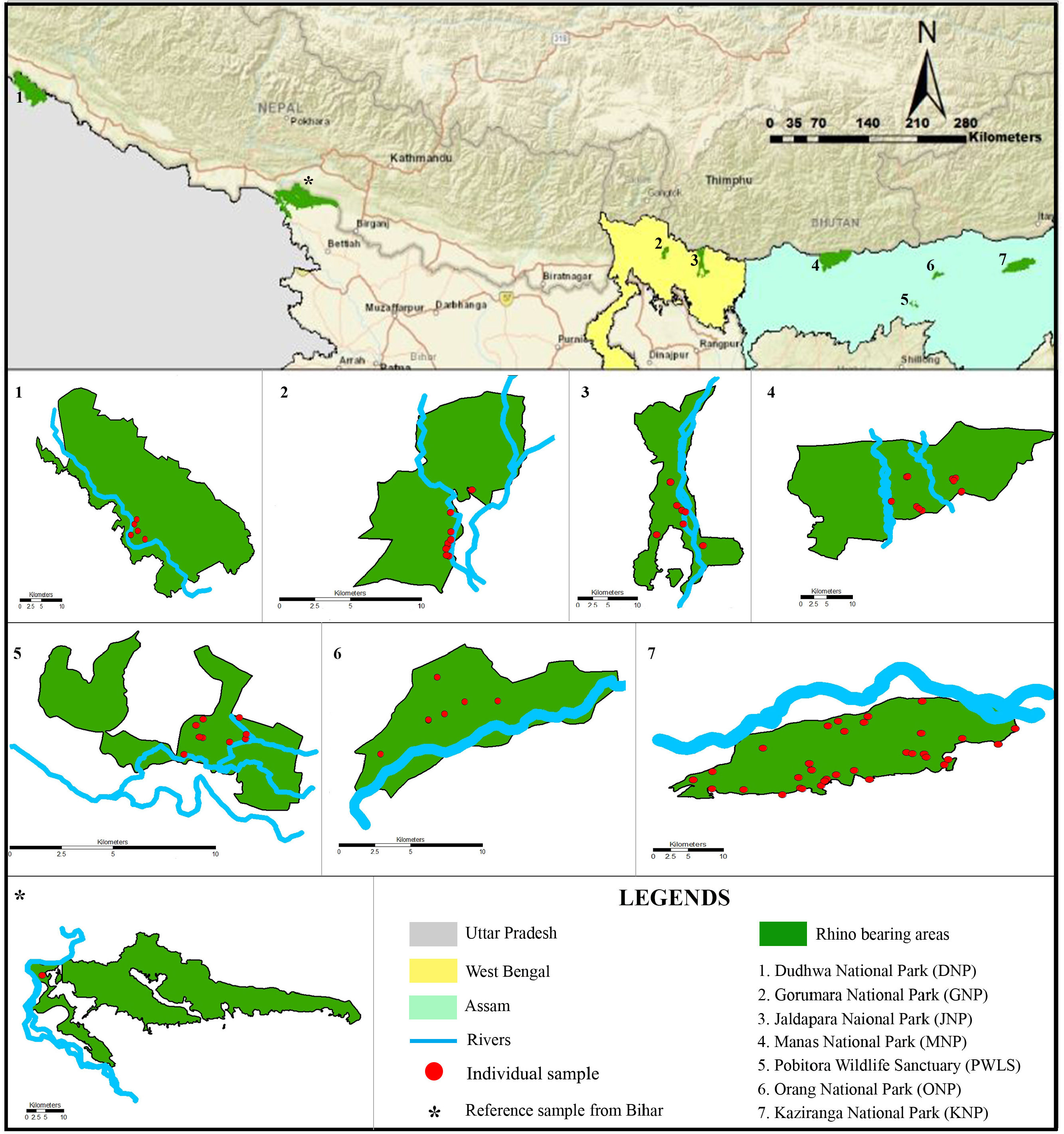
Map of the study area and distribution of the final samples used in this study (n=111). The top plate shows the position of the rhino-bearing parks across three Indian states (Uttar Pradesh, West Bengal and Assam). The reference sample of wild rhino received from the state of Bihar is also presented.

### 2.3 Laboratory work

#### 2.3.1 DNA extraction

Tissue DNA was extracted using already established protocol for Indian rhino mentioned in Ghosh et al., (2021). For dung samples, a modified protocol from Biswas et al., (2019) was used. In brief, samples were digested overnight with a combination of 700 μl ATL and 65 μl Proteinase K (20mg/ml) at 56◦C, followed by QIAamp DNA Tissue Kit (QIAGEN Inc., Hilden, Germany) protocol with adjusted volumes. DNA was eluted twice in 100 μl preheated (70 ◦C) 1X TE buffer and stored in -20°C freezer. Extraction negative was used for each set of extraction (n=23) to monitor possible contamination.

#### 2.3.2 PCR amplification and sequencing

To assess genetic variation of the extant rhino populations, complete mitogenome data was generated for representative samples from each park (n=15, see Table A1 for details) and one from the Valmiki National Park (Bihar Forest Department). Mitogenome sequencing was performed using published 23 overlapping primers (Hassanin et al., 2009). For annealing temperature standardisation, gradient PCR was set in 10 μl reactions containing 4 μl of 2X Qiagen PCR buffer mix (QIAGEN Inc., Hilden, Germany), 1 μl of primer (3 μM), 2 μM BSA (4 mg/ml), 1.4 μl of RNase free water and 5 ng of rhino tissue DNA. PCR conditions included an initial denaturation (95 °C for 15 min); 35 cycles of denaturation (95 °C for 30 s), annealing (50-60 °C gradients for 40 s) and extension (72 °C for 40 s); followed by a final extension (72 °C for 10 min). During each set of reactions, PCR and extraction negatives were included to monitor contamination. Amplified products were visualized with 2% agarose gel, cleaned with Exonuclease (Thermo Scientific, Waltham, USA) and Shrimp Alkaline Phosphatase (Amresco, Solon, USA) mixture and sequenced bidirectionally in an ABI 3500XL bioanalyzer (Applied Biosystems). Out of these 23 primers, two did not show amplification in any samples. The remaining sequences (n=21 from 16 individuals) were aligned with the available one-horned rhino mitogenome (Genbank: X97336, Xu et al., 1996) in Mega v7 (Kumar et al., 2016). Primers were designed manually in the flanking conserved regions adjacent to the gaps and sequences were generated from all the samples (n=16).

The complete mitochondrial sequences (n=16) were aligned and manually screened to identify the segregating sites. Further, a total of 15 primers were designed to amplify all the polymorphic sites as <500 bp fragments to ensure higher success rate from fresh dung DNA. These primers were standardised following same protocol described above. For all field collected samples (tissue= 58 and dung=88) individual identification was performed using a panel of 14 microsatellites (described in Ghosh et al., 2021). After PCR amplification and genotyping of the markers, samples with <12 loci data were removed from downstream analysis. Further, genetic recaptures were removed and to ensure uniform representation we selected one sample from every area with clustered individuals. Sequence data (2531bp covering seven genes) was generated for the selected individuals to assess phylogeography patterns.

### 2.4 Analysis

#### 2.4.1 Complete mitogenome annotation and comparative analysis

All rhino sequences (n=16) were aligned in Mega v7 to generate a complete mitogenome sequence and manually checked to identify any nucleotide ambiguities. Annotation was done using MITOS2 web (Bernt et al., 2013) and mitogenome map was created with OGDRAW (Greiner et al., 2019). To ascertain species-wise mitochondrial DNA diversity these sequences were aligned with already published rhino mitogenome sequences from *Diceros bicornis* (n=2, Genbank: FJ905814, NC012682 (Willerslev et al., 2009)), *Ceratotherium simum* (n=2, Genbank: Y07726, NC001808 (Xu et al., 1997)), *Dicerorhinus sumatrensis* (n= 15, Genbank: MF066629-MFO66643 (Steiner et al., 2017) and *Rhinoceros sondaicus* (n=6, Genbank: FJ905815 (Willerslev et al., 2009), MK909142, MK909146, MK909148, MK909149, MK909151 (Margaryan et al., 2020)). We calculated number of segregating sites (S), nucleotide (π) and haplotype diversity (Hd) using DnaSP v.5 (Librado and Rozas, 2009) for all genes in the mitogenome.

#### 2.4.2 Genetic diversity in Indian rhinos

Population-wise basic indices of genetic variations (S, π and Hd) were calculated for concatenated sequence data (2531bp from seven genes) using DnaSP v.5 followed by a median joining (Bandelt et al., 1999) haplotype network constructed in PopART v. 1.7 (Leigh and Bryant, 2015). To ascertain any possible population structure a Bayesian approach implemented in BAPS v.5.3 was used as it considers linked loci data (Corander and Tang, 2007). Pairwise F_st_ and differential hierarchical AMOVA analysis was performed using Arlequin v. 3.0 (Excoffier et al., 2005) to confirm the pattern found in BAPS analysis.

#### 2.4.3 Estimation of clade-specific divergence times

To identify the clades, Bayesian phylogeny was constructed with MrBayes v. 3.2.7 (Huelsenbeck and Ronquist, 2001) using 16 Indian rhino mitogenome and Javan rhino sequence (outgroup, as they are the sister clade of one-horned rhinoceros) (Harley et al., 2016). Analysis was conducted using GTR+G substitution model determined by jModelTest v2.1.3 (Darriba et al., 2012) (based on Akaike Information Criteria). The MCMC parameters included 2 runs of four chains each of 15 million generations with sampling after 1000 generations till split frequencies were below 0.01. Posterior probabilities were calculated for each node.

To estimate divergence among clades, rate of mutation for Indian rhino was calculated using BEAST v.2.3.6 (Drummond et al., 2007). Analysis was performed with five extant rhino mitogenome (without D-loop) (n=11 sequences, India=7, Java=1, Sumatra=1, White=1, Black=1) along with horse (*Equus caballus,* Genbank: NC001640), donkey (*Equus asinus,* Genbank: NC001788), Asiatic wild ass (*Equus hemionus*, Genbank: NC016061) and zebra (*Equus zebra*, Genbank: NC018780) as outgroups. GTR+G substitution model was selected through jModelTest v2.1.3 for this multi-species data. Birth-death speciation was considered as tree prior (Ritchie at al., 2017; Steiner et al., 2017) along with uncorrelated relaxed log normal clock (Drummond et al., 2006; Steiner et al., 2017). During analysis, four established internal node calibration points (based on fossil records) with normal distribution priors were employed: (i) Caballine split (4± 0.5 million years ago (Mya)) (Eisenmann, 1992; Orlando et al., 2013); (ii) late Oligocene diversification of rhino groups (26± 3.5 Mya) (Hooijer, 1946); (iii) split of rhinoceros genus (3± 0.5 Mya) (Liu et al., 2021); (iv) origin of the perissodactyls (55± 3 Mya) (Prothero and Schoch, 1989; Benton and Donoghue, 2007). The first three calibration points were considered as monophyletic constraint (Drummond et al., 2006) as the last point includes both ingroup and outgroup taxa.

tMRCA (the Most Recent Common Ancestor) was inferred using the estimated mutation rate with lognormal distribution under strict molecular clock (intra species data, n=16) (Yoder and Yang, 2000; Ritchie et al., 2017). MCMC runs included 100 million generations, sampled at every 10,000 states with 10% burn-in. Data convergence was checked with Tracer v. 1.5 (Rambaut and Drummond, 2007) and the final tree (with maximum clade credibility) was estimated with TreeAnnotator (Helfrich et al., 2018) and visualised using FigTree v.1.4.2 (Rambaut, 2009).

## 3. Results

### 3.1 Rhino mitogenome data and comparative analyses

Sequencing with 23 primers (Table A2) generated 16828bp mitogenome (Figure A1) for wild Indian rhino (n=16, Genbank: MZ736693-MZ736708). Composition analysis revealed AT-skewed mitogenome with 13 protein coding genes, 22 tRNA, 2 ribosomal genes and a non-coding control region (Table A3). Comparative analyses with other rhino species (Figure A2) revealed that the Indian rhinos have low segregating sites (S_Java_=15514, S_Africa_=10680, S_Sumatra_=130, S_India_=18) and nucleotide diversity (π_Java_=0.56, π_Africa_=0.43, π_Sumatra_=0.003, π_India_=0.0005) but high haplotype diversity (Hd_Sumatra_=0.96, Hd_India_=0.93, Hd_Java_=0.91, Hd_Africa_=0.67). Both African rhino species (white and black rhino) data were combined for this analyses as no intra-species variation was observed in the available data.

### 3.2 Phylogeography of wild Indian rhinos

Out of 15 primers designed to assess genetic variation, eight were finally used (Table A2) to amplify all 22 polymorphic sites (covering 2531bp sequence) of rhino mitogenome. This data was generated for additional 95 unique individuals (n= 56 tissue and 39 dung) (Genbank: MZ771364-MZ771458, MZ771459-MZ771553, MZ771554-MZ771648, MZ771649-MZ771743, MZ771744-MZ771838, MZ771839-MZ771933 and MZ771934-MZ772028). Remaining samples could not be used as they were genetic recaptures (n=13), individuals representing close geographic clusters (n=24) or low amplification success (n=12). Median joining network (n=111 individuals) showed a total of 30 haplotypes (h) across India. Majority of these haplotypes (93.3%, n=28) were from Assam whereas both West Bengal (one haplotype) and Uttar Pradesh (one haplotype) populations were found to be monomorphic (Fig. 2). The sequence from Bihar rhino individual was identical to the Uttar Pradesh population. Population-wise genetic variation indices (Table 1) showed overall highest values for KNP (n= 46; S=18, h=19, π=0.0021, Hd= 0.85), followed by MNP (n=12; S=14, h=6, π=0.0023, Hd= 0.89), ONP (n=12; S=9, h=6, π=0.0016, Hd= 0.89) and PWLS (10; S=2, h=3, π=0.0002, Hd= 0.51). Bayesian genetic clustering corroborated with the earlier pattern where samples from West Bengal and Uttar Pradesh formed distinct clusters whereas Assam showed geographically intermixed haplogroups (Fig. 2). The genetic differentiation (pairwise F_st_) values among these three clusters were significantly high ranging from 0.68-1 (Table 2, indicating highly structured populations). The hierarchical AMOVA analysis using two separate groupings: a) seven populations and b) three states showed higher within population (50%) and between group variance (45%) also indicating strong genetic structures in Indian rhinos (Table 2).

**Figure 2:**
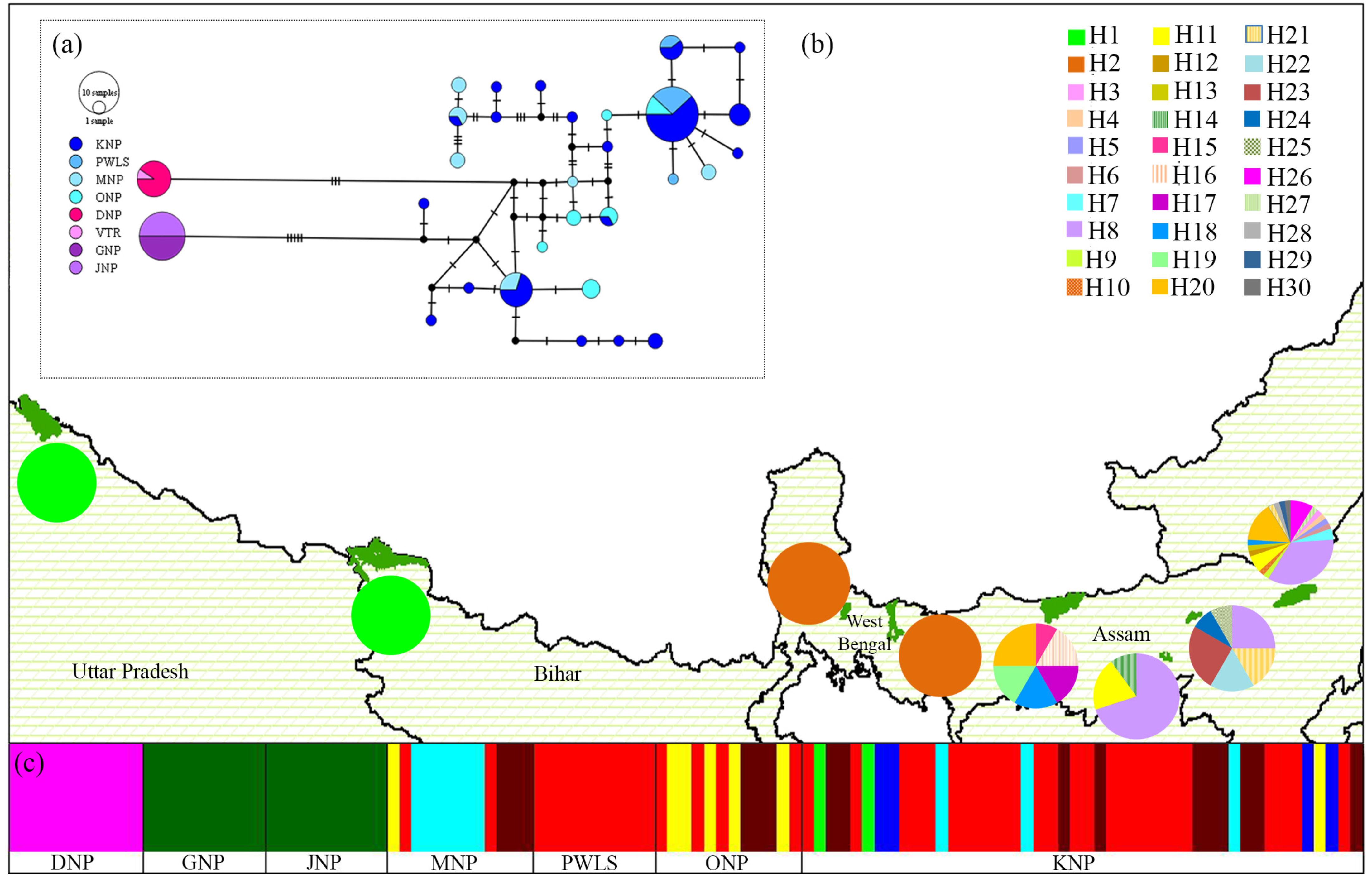
Representation of mtDNA variations and genetic structure in Indian rhinos based on 2531 bp concatenated sequence covering all polymorphic sites across seven genes. (a) Median joining network with park-level colour codes; (b) Haplotype frequencies at each of the sampled areas covering all variations (n=30 haplotypes); (c) Bayesian clustering shows monomorphism in Uttar Pradesh (with sample from Bihar, n=11) and West Bengal (n=20) populations and polymorphism in Assam (n=80).

**Table 1:** mtDNA diversity indices of all seven rhino populations in India (n=110).

| Protected areas | Sample size | Segregating sites (S) | Haplotypes (h) | Haplotype diversity (Hd) | Nucleotide diversity ($\pi$) |
| --- | --- | --- | --- | --- | --- |
| Kaziranga National Park, Assam | 46 | 18 | 19 | 0.85 | 0.0021 |
| Pobitora Wildlife Sanctuary, Assam | 10 | 2 | 3 | 0.51 | 0.0002 |
| Manas National Park, Assam | 12 | 14 | 6 | 0.89 | 0.0023 |
| Orang National Park, Assam | 12 | 9 | 6 | 0.87 | 0.0016 |
| Dudhwa National Park, Uttar Pradesh | 10 | 0 | 1 | 0 | 0 |
| Gorumara National Park, West Bengal | 10 | 0 | 1 | 0 | 0 |
| Jaldapara National Park, West Bengal | 10 | 0 | 1 | 0 | 0 |
| **Total** | **110** | **22** | **30** | **0.89** | **0.0028** |

**Table 2:**
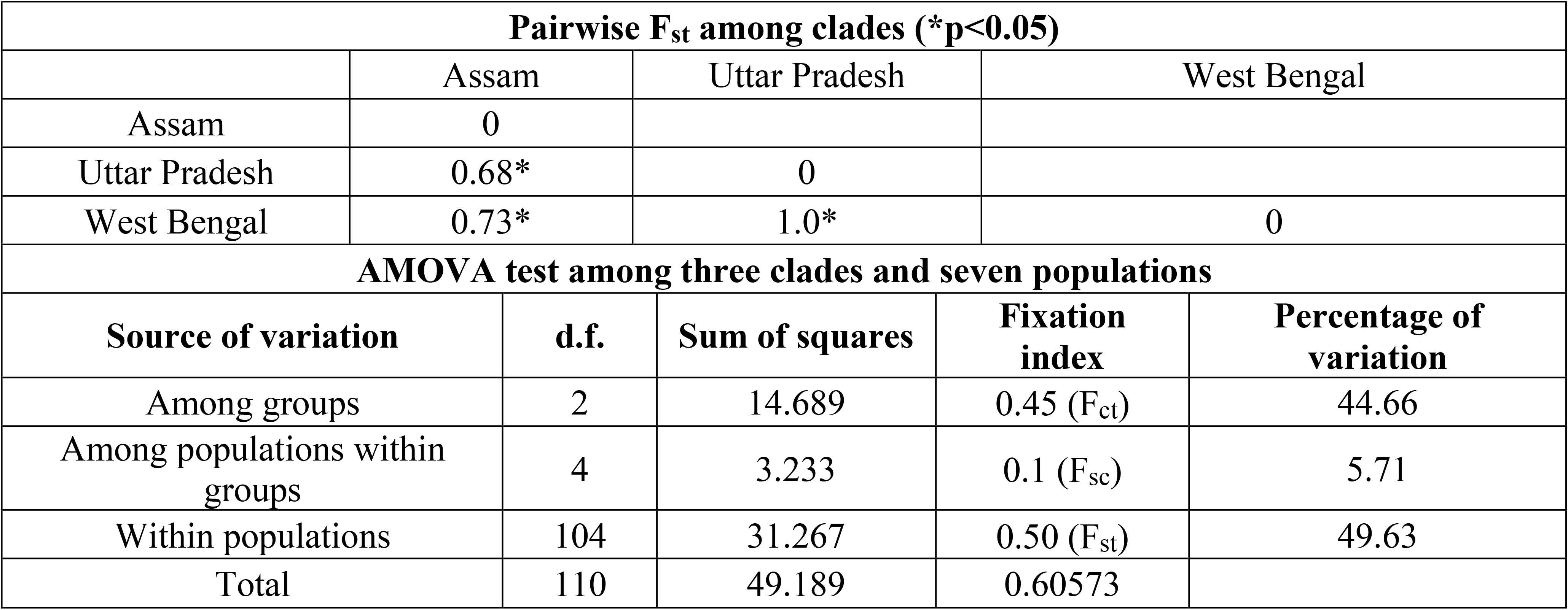
Results of Pairwise genetic differentiation and hierarchical AMOVA test (Bihar sample considered under Uttar Pradesh clade).

### 3.3 Divergence time of different rhino clades

The Bayesian phylogeny showed similar pattern of three clades: the West Bengal samples representing basal clade (Fig. 3, node G) followed by Assam (nodes C-E) and Uttar Pradesh (along with the Bihar sample) representing two sister clades (node F). Based on the calibrated root nodes and Indian rhino-specific mutation rate (1.2 x 10^-4^ mean rate of substitution per site per million years, Figure A3), tMRCA analysis suggested a divergence period spanning from 0.95 (HPD 1.36-0.81 Mya) to 0.05 (0.15-0.01 Mya) Mya (Fig. 3). Our results indicated the divergence of Indian rhinos ∼ 0.95 Mya (node A, Fig. 3) corresponding to the emergence period of modern mammalian ancestors in the subcontinent (Lal, 2016; Patnaik, 2016). Next, the Assam population separated ∼0.5 Mya (HPD 0.68-0.33 Mya, nodes B & C, Fig. 3) from the remaining clades. This is supported by reports of multiple rhino movements away from Assam (along Siwalik as well to Siva-Malayan region) during this period (van den Bergh et al., 2001; Patnaik, 2016). At population level, results suggest a relatively older coalescence of Assam ∼0.19 Mya (HPD 0.3-0.07 Mya, node D & E, Fig. 3) compared to West Bengal and Uttar Pradesh (∼0.05 Mya, HPD 0.15-0.01 Mya, node F & G, Fig. 3). This period (0.12-0.01 Mya) is known for confinement of rhinoceros to the north and north-east of India due to monsoon intensification and grassland dominance (Patnaik, 2013, 2016; Lal, 2016).

**Figure 3:**
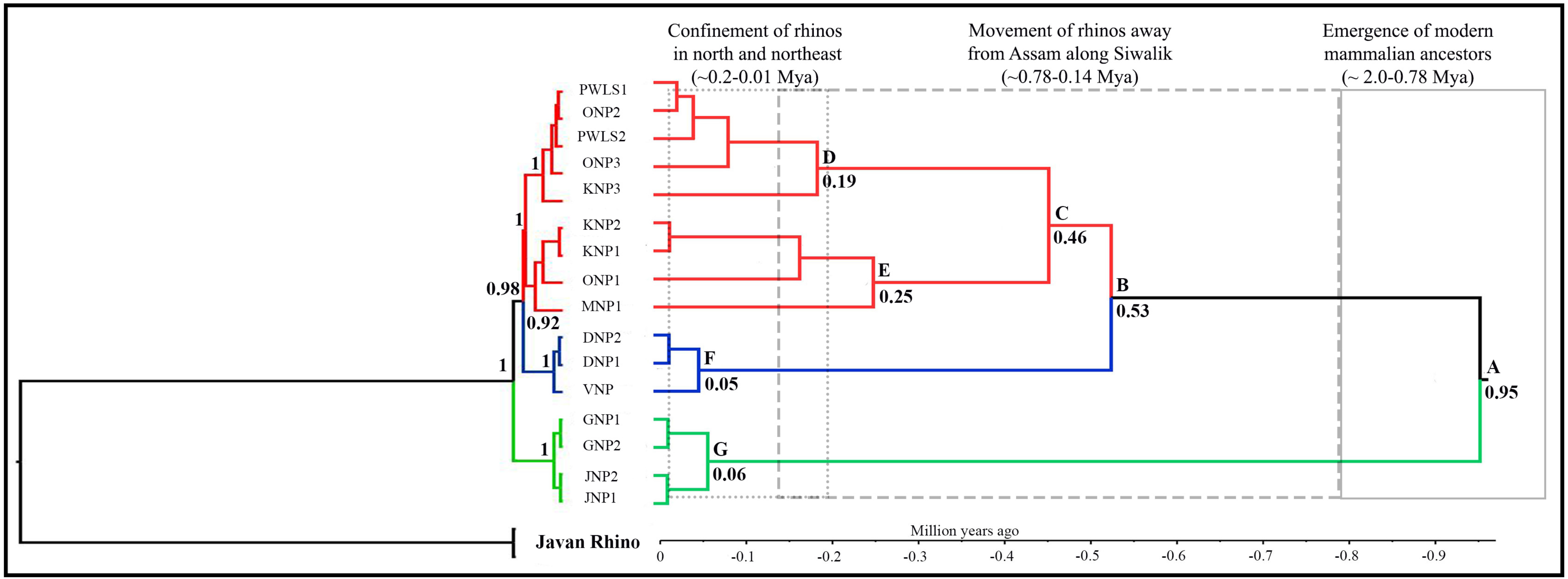
Phylogenetic relationship and assessment of divergence time in Indian rhino populations. The left pane shows the basal clade of West Bengal samples (green) and sister clades representing Uttar Pradesh (blue) and Assam (red). Javan rhino sequence was used as outgroup. The posterior probability values (≥0.9) are shown in bold. The right pane indicates the divergence of Indian rhinos, where the Assam population diverged first (∼0.5 Mya), followed by parallel divergence of West Bengal and Uttar Pradesh (0.06-0.05 Mya). Node-specific ages are marked (with posterior probability values ≥0.9). The major corroborating paleobiogeographical events are presented above.

## 4. Discussion

This study presents the most extensive mitochondrial DNA phylogeography of one-horned rhinos across its Indian distribution. Careful considerations involving mitogenome sequencing of representative samples across Indian rhino-bearing areas, identification of all polymorphic regions and their amplification from spatially-covered rhino samples helped us achieving accurate assessment of mtDNA variations. To the best of our knowledge, this is the first report of wild Indian one-horned rhino mitogenome from all the extant populations. Despite relatively similar haplotype diversity of Asian rhinos (India-0.93 (16 samples), Sumatra-0.96 (15 samples), and Java-0.91 (6 samples), respectively), Indian rhino mitogenome showed much lower values for segregating sites and nucleotide diversity. Such mitogenome comparisons may be affected by limited sample size (in African rhinos, Moodley et al., 2017, 2018) or representation of historical genetic variations (in Javan rhinos, Margaryan et al., 2020). However, it was surprising to observe that despite similar historical demographic incidences (severe population decline due to habitat shrinkage (Patnaik, 2016; Mays et al., 2018) and anthropogenic pressures (Das et al., 2015; Steiner et al., 2017) Indian rhino retain much lower genetic variation than their Sumatran counterpart. This can be potentially attributed to recovery of the Indian species from extremely low founder population (as indicated by high Hd but low π) (Avise, 2000; da Silva et al., 2018).

As expected, the phylogeography data (2531bp mtDNA, n= 16 samples) revealed higher number of haplotypes than the mitogenome data (n=30 haplotypes) (due to large sample size). The only other study on one-horned rhino mtDNA variations (based on partial control region sequences, 428 bp) reported 10 haplotypes (Kaziranga National Park, India-4 and Chitwan National Park, Nepal-6, respectively) and moderate level of genetic difference (F_st_ value of 0.39 between them) (Zschokke et al., 2011). Careful scrutiny of our data revealed that all the polymorphic sites (or identified segregating sites) were found in fixed positions within one-horned rhino mitogenome (Table A4) across India. Given the fixed positions of these sites (and no new variations in the samples) and our sampling coverage it can be presumed with high confidence that all potential rhino mtDNA haplotypes from India have been reported in this study. This claim is also supported by the similar haplotype diversity values from the mitogenome and the phylogeography datasets (0.93 and 0.9, respectively). Our study also shows that the Indian rhinos have the highest number of haplotypes compared to the other species/subspecies reported so far (Steiner et al., 2017, Moodley et al., 2017, 2018; Margaryan et al., 2020). The clustering analysis of the concatenated rhino sequences showed three distinct genetic clades (corresponding to the states of Assam, West Bengal and Uttar Pradesh) with high F_st_ value (0.68-1), corroborating with the haplotype network patterns. Mantel test (−0.83, p=1) confirmed that such strong genetic structuring is not due to isolation by distance pattern, but driven by lineage-specific evolutionary history (as suggested by AMOVA results). Such pattern of higher within population and between group variance (50% and 45% in Indian rhinos, respectively) has also been described in other species (barking deer- Martin et al., 2017, dog- Pang et al., 2009). Interestingly, we found that the sequence from the Bihar sample (representing samples from Nepal) was identical to the Uttar Pradesh sequences, including the state-specific SNPs. Though surprising, this pattern was expected as the founder animals of the reintroduced Uttar Pradesh population were sourced from Chitwan National Park, Nepal (four dominant breeding females) and Pobitora Wildlife Sanctuary of Assam (dominant breeding male) (Talukdar et al., 2012). Further comparison of 13 partial D-loop sequences from Chitwan National Park, Nepal (Zschokke et al., 2011) confirmed this pattern, indicating that the mtDNA signature of the Uttar Pradesh population belongs to Nepal. Given that the entire Uttar Pradesh rhino population showed only one haplotype, future studies need to evaluate the mtDNA variation in the Nepal population.

The phylogenetic analyses reconfirmed the relationship among the existing members of the Rhinocerotidae family (Xu et al., 1996; Steiner et al., 2011, 2017) and the genetic clades (similar to the phylogeographic pattern) within India. In the analyses, we used only the existing rhino species/subspecies data (Woolly rhinoceros sequence was not used) and the Sumatran and African rhino formed sister clades, separated from the Indian clade. However, within these reciprocal monophyletic clades in India the Assam population diverged first (0.5 Mya, Fig. 3) followed by the other two groups (West Bengal and Uttar Pradesh/Nepal), diverging around similar time period (0.06-0.05 Mya, Fig. 3). The parallel divergence of the West Bengal and Uttar Pradesh is also supported by no shared genetic signatures between them (Fig. 2). The molecular dates were comparable to other published literature on rhino evolution (Lal, 2013; Patnaik, 2013, 2016) and supported by the paleobiogeographic history of the Indian subcontinent (Patnaik, 2013, 2016). For instance, the inward movement of rhinos from Assam along Siwalik (0.68-0.33 Mya, node B & C) coincides with drop in the sea level which facilitated movement of multiple genera (for example, *Elephus, Panthera*, *Rhino*, *Muntiacus* etc.) through Siva-Malayan route (van den Bergh et al., 2001; Vidya, 2015). Report of one-horned and Javan rhino co-existence in Bhutan ∼0.56 Mya (Margaryan et al., 2020) provide further support of such movements. Finally, the coalescence time of the Indian clades corresponds to Holocene climatic optimum period known for monsoon intensification in north and north-east part of India resulting in range contraction for grassland dependent species (Lal, 2013; Patnaik, 2016; Kumar et al., 2017). We feel that our approach of using taxon-specific mutation rate and fossil data for node calibration has resulted in achieving such meaningful estimates of tMRCA. Future efforts should try to include molecular data from historical/ancient samples to tighten the variance associated with divergence estimates (Drummond et al., 2007). Overall, this approach reiterates the critical importance of large datasets (whole mitogenome from multiple individuals in this case), informative prior settings and its assessment with posterior outputs, taxon-specific mutation rate, node calibration points etc. for accurate tMRCA estimation (Yoder and Yang, 2000; Drummond et al., 2006; Subramanian et al., 2009; Knaus et al., 2011; Ritchie at al., 2017).

The spatially exhaustive sampling coverage and the patterns of population structure brings out some critical conservation perspectives for the Indian rhinos, particularly when compared with the other Asian counterparts. For example, the different divergence time and high genetic differentiation among the three populations of wild rhinos warrant a consideration of possible subspecies designations for them. When looked carefully at the Sumatran rhino evolution, the three subspecies (designated based on morphological differences, Groves, 1967) shows very similar kind of divergence time history (Steiner et al., 2017) and much less genetic differentiation (0.098) than the Indian rhinos, providing strong argument for consideration of genetic subspecies for the three Indian populations. However, these populations are morphologically undistinguishable and interbreed among themselves (Dudhwa, Uttar Pradesh population is genetically mixed, Talukdar et al., 2012), and therefore would be meaningful to be recognised as ‘Evolutionary Significant Units (ESUs)’ as the current information emphasize their evolutionary histories. Based on the data, it is also important to point out that the conservation and management efforts need to equally focus for each of these populations and consider detailed understanding of their history. For example, the Uttar Pradesh and West Bengal population show state-specific monomorphic haplotypes representing unique but genetically depauperate populations. However, ever-continuing anthropogenic pressures and loss of habitat is further increasing the effects of fragmentation as seen in West Bengal where recent study showed strong genetic structure between the existing rhino populations (F_st_ value of 0.31 based on 11 microsatellite data) (Das et al., 2015). On the other hand, the reintroduced rhino population in Assam (Manas National Park) showed much higher mtDNA variation (six haplotypes), possibly due to periodic supplementation of individuals of varied genetic ancestry across different wild rhino populations (Talukdar et al., 2020). Careful management decisions in terms of selecting genetically variable founder animals, multiple reintroduction events etc. would be critical for future rhino reintroduction programs in India (Weeks et al., 2015; Jhala et al., 2021). Given that multiple reintroduction programs are planned as per the ‘National Conservation Strategy for the Indian One horned rhinoceros (*Rhinoceros unicornis*), Government of India, Ministry of Environment Forest and Climate Change, 2021’ objectives (in the states of Uttar Pradesh, Bihar, West Bengal and Assam) in near future, the genetic signatures described in this study would be very helpful in selecting appropriate areas to identify founder animals.

## 5. Conclusion

The one-horned rhino was found throughout the Indo-Gangetic plains during the early 20^th^ century (Rookmaaker, 1983; Rookmaaker et al., 2016) but faced drastic reductions in distribution and population size (including local extinctions) (Talukdar et al., 2008; Pant et al., 2019) and shown high extinction probability among the Indian megaherbivores (Karanth et al. 2010). However, strong conservation and management initiatives since late 1990s resulted in one of the most successful species recovery (increase in population size) in wild across the world (Menon, 1996; Rookmaaker et al., 2016; Pant et al., 2019). Despite a relatively long history of effective management efforts, information on genetic status of the species has not been used in translocation programmes (possibly due to lack of sufficient data). Rhino is a highly conservation-dependent species and any future translocation without inputs from the genetic data from the source populations may further impact the genetic status of the new habitats (as reported in case of Florida panther - Hostetler et al., 2010, Johnson et al., 2010; Cheetah- O’Brien et al., 2017; Gharial- Sharma et al. 2021). We present the first assessment of range-wide mitogenome diversity in Indian rhinos where we emphasize the importance of large data, spatial sampling coverage of populations and evolutionary history as fundamental information for future population reintroduction/ recovery programs. Our results are important for Indian rhino conservation because they suggest higher genetic diversity than earlier reported (Zschokke et al., 2011). However, the existing habitats are small, disjunct, isolated and reaching their respective carrying capacities (Pant et al., 2019; Jhala et al., 2021) and conservation options are becoming limited except establishing new habitats and translocation-driven population enhancement (Jhala et al., 2021). We believe that the genetic information provided here will assist in identifying appropriate source populations and maintain adequate genetic diversity in the existing (and new) rhino populations, thereby ensuring evolutionary, ecological and demographic stability for their future survival.

## Acknowledgment

We thank the Ministry of Environment, Forest and Climate Change, Government of India for all support to implement this project. We thank Forest Departments of Assam, West Bengal and Uttar Pradesh for necessary permits and field support during sampling. We thank the WWF-India team for their field and logistic support. We appreciate the help from Mirza Ghazanfarullah Ghazi regarding marker aliquots during initial standardizations. Our thanks to Ankit Pacha, Bhim Singh, Surya P. Sharma and Dr. Kunal Arekar for their technical inputs in analysis. We thank the Director, Dean, Research Coordinator and Nodal Officer of the Wildlife Forensics and Conservation Genetics Cell for their support. Ministry of Environment, Forest and Climate Change, Government of India and WWF-India funded the work.

## CRediT author statement

**Tista Ghosh:** Conceptualization, Methodology, Investigation, Validation, Formal Analysis, Data Curation, Writing-Original draft and editing, Visualisation

**Shrewshree Kumar:** Investigation, Validation, Formal Analysis, Data Curation, Writing-review and editing, Visualisation

**Kirtika Sharma:** Investigation, Validation, Data Curation, Writing-review

**Parikshit Kakati:** Resources, Methodology, Project administration, Writing-review

**Amit Sharma:** Resources, Project administration, Funding acquisition, Writing-review

**Samrat Mondol:** Conceptualization, Resources, Writing-Original draft and editing, Project administration, Funding acquisition, Supervision

---

**Supplementary Fig.A1:**
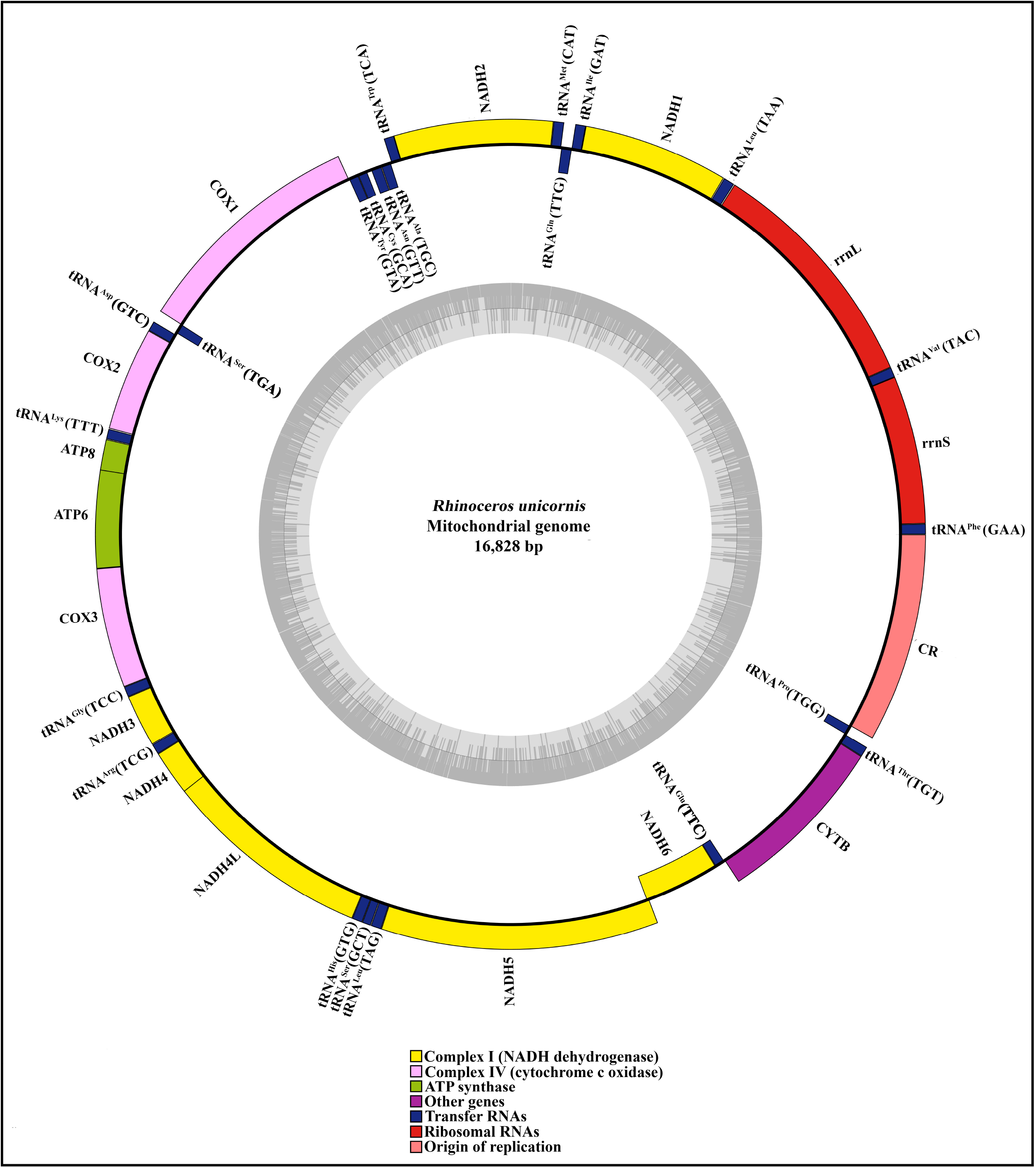
Whole mitogenome organisation and annotation of *Rhinoceros unicornis*.

**Supplementary Fig.A2:**
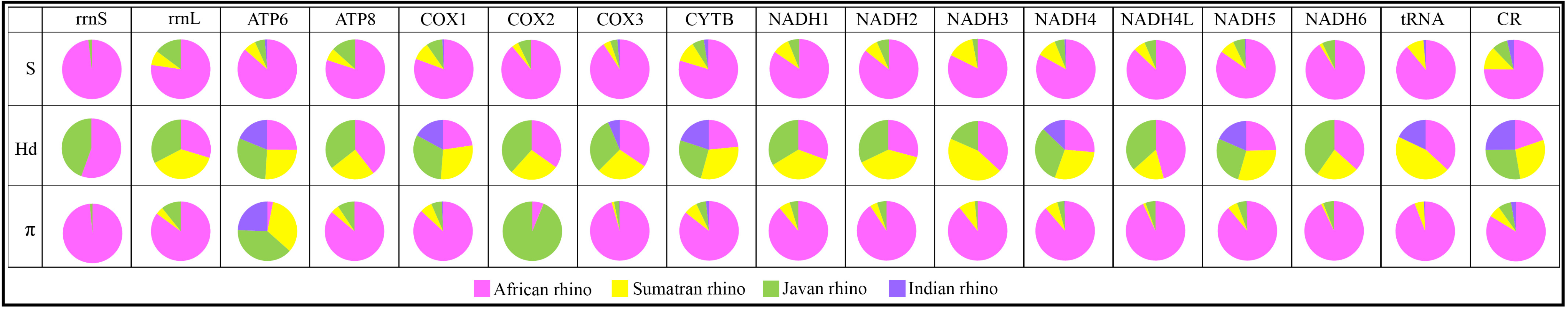
Gene-wise comparative analyses of polymorphism indices (S, Hd and π) for all extant rhino species.

**Supplementary Fig.A3:**
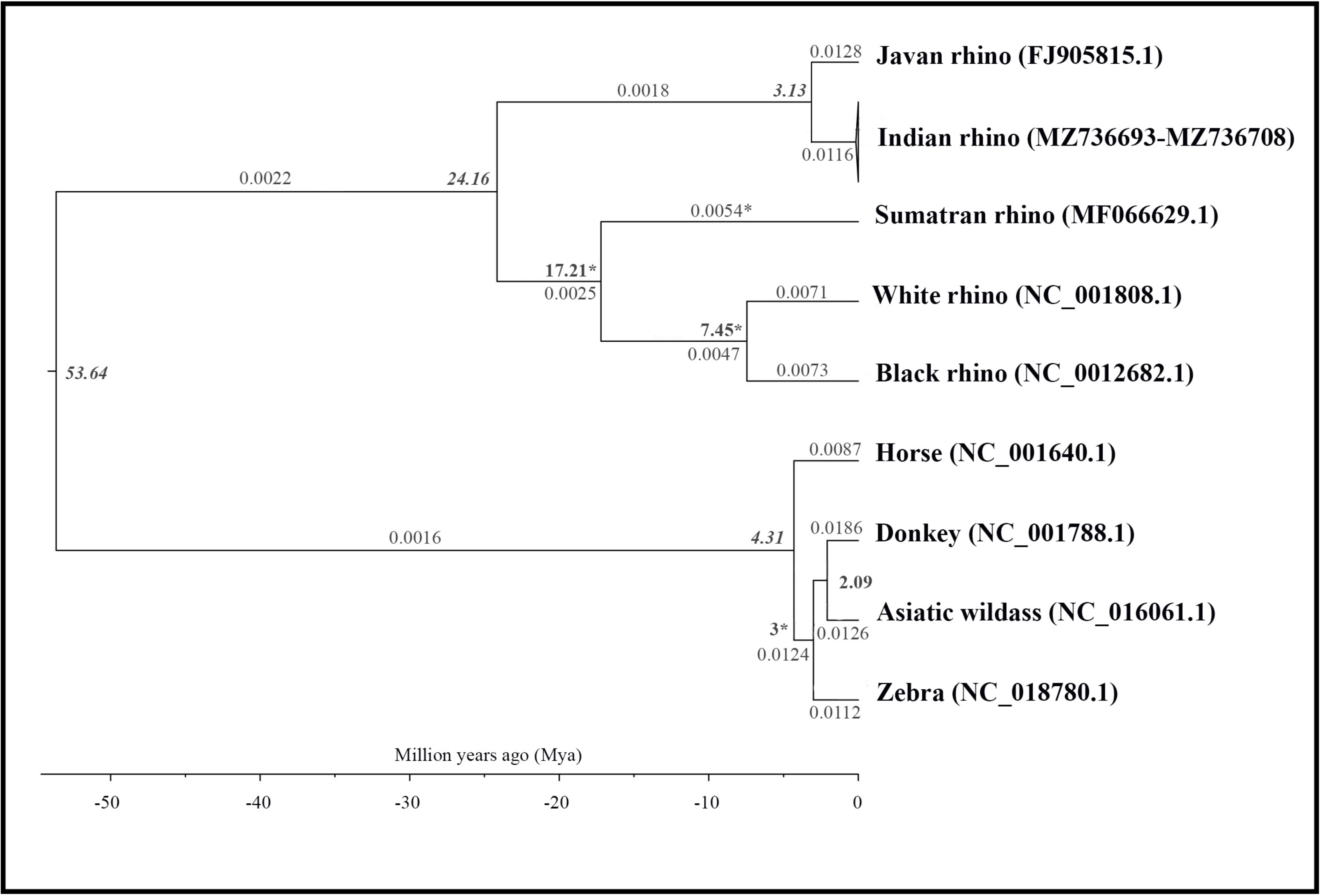
Estimation of mitogenome mutation rate for Indian rhino using Caballine (Zebra, Donkey, Asiatic Wild Ass and Horse) as outgroup. Internal node calibration points are presented in italics (bold). Estimated node ages and branch mutation rates with available reference in literature is marked with * (Fernando et al., 2006; Steiner et al., 2011; Steiner et al., 2017; Liu et al., 2021).

**Supplementary Table A1:**
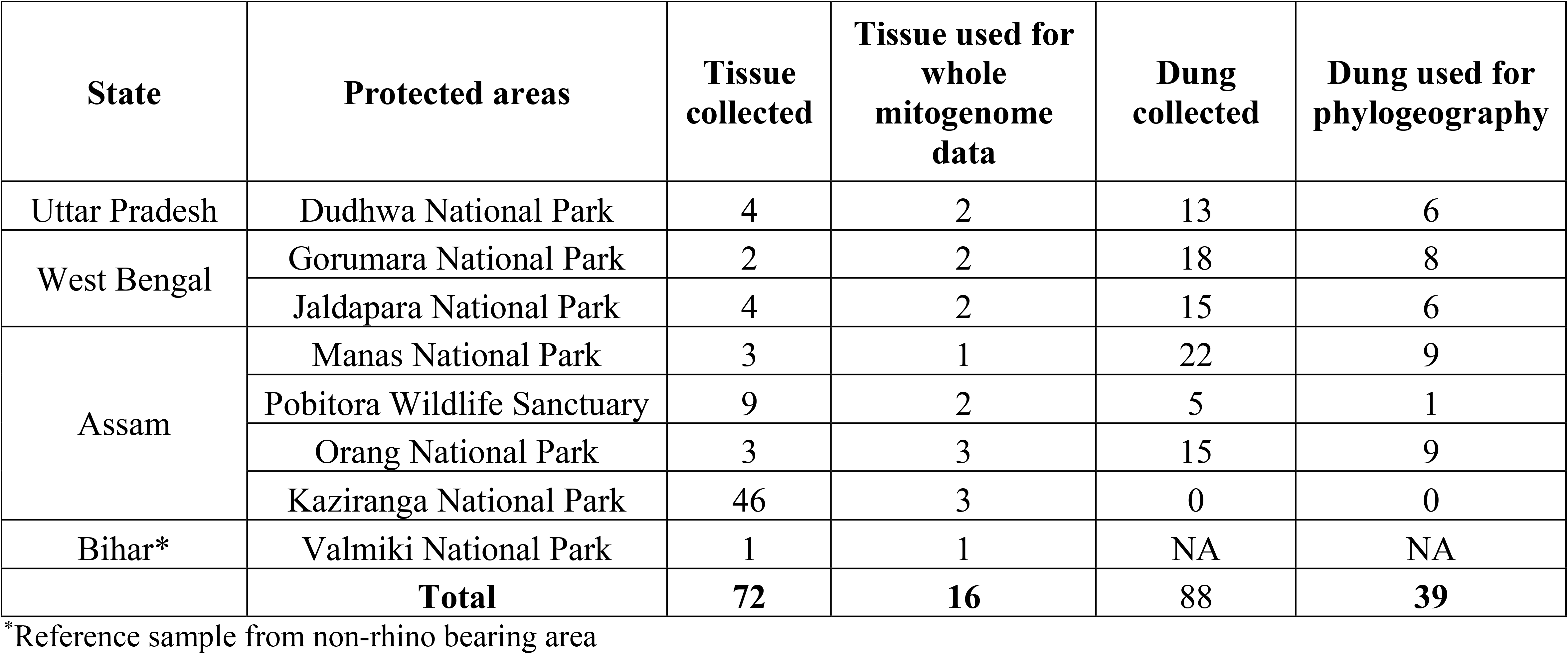
Sample (both tissue and dung) and data details of Indian rhinos used in this study.

**Supplementary Table A2:**
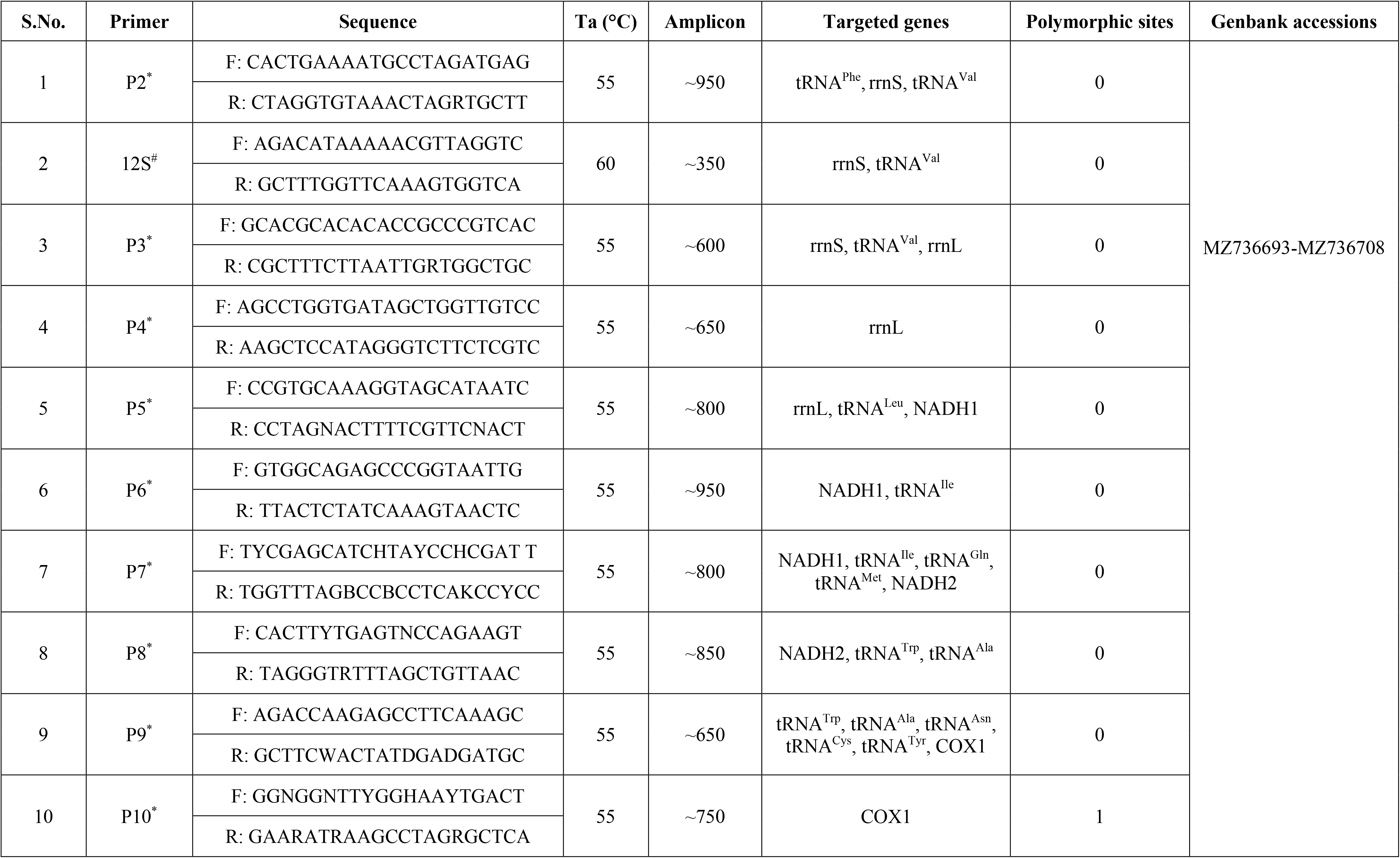
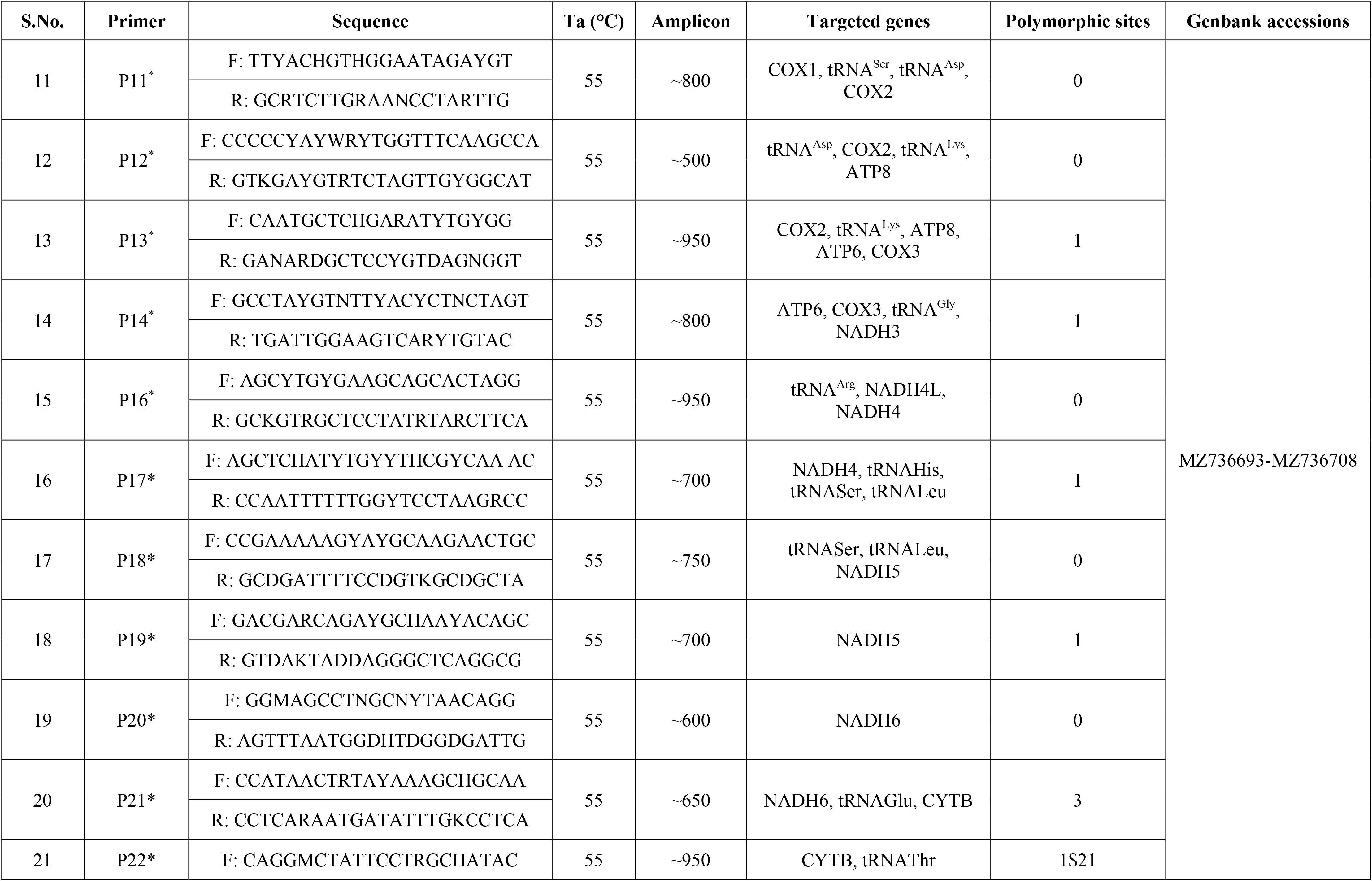
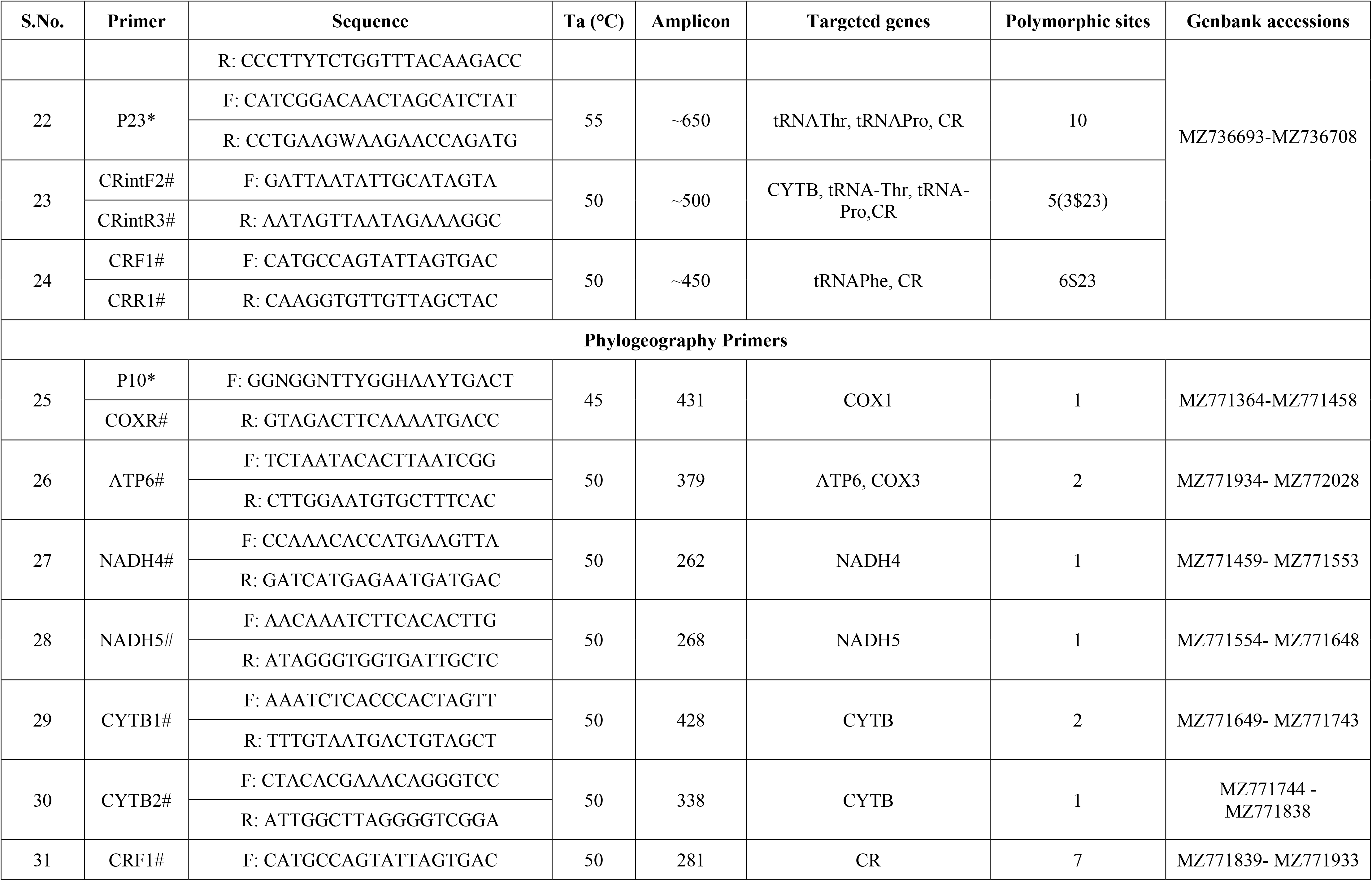
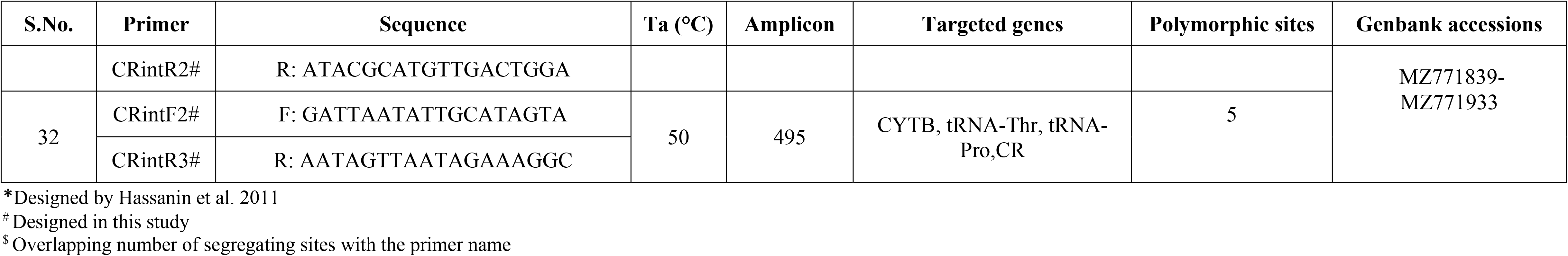
Details of the primers used in sequencing Indian rhino whole mitogenome and phylogeography data.

**Supplementary Table A3:** Mitogenome organization in *Rhinoceros unicornis*. Codons respective to each tRNA are mentioned in parenthesis.

| Genes | Position |  | Gene length | Spaces/<br>overlap | Strand | Codons |  |
| --- | --- | --- | --- | --- | --- | --- | --- |
|  | Start | Stop |  |  |  | Start | Stop |
| tRNA<sup>Phe</sup> (GAA) | 1 | 69 | 69 | 0 | H |  |  |
| rrnS | 70 | 1041 | 972 | -1 | H |  |  |
| tRNA<sup>Val</sup> (TAC) | 1041 | 1107 | 67 | 0 | H |  |  |
| rrnL | 1108 | 2681 | 1574 | 3 | H |  |  |
| tRNA<sup>Leu</sup> (TAA) | 2685 | 2759 | 75 | 2 | H |  |  |
| NADH1 | 2762 | 3718 | 957 | -1 | H | ATG | TAA |
| tRNA<sup>Ile</sup> (GAT) | 3718 | 3786 | 69 | -3 | H |  |  |
| tRNA<sup>Gln</sup> (TTG) | 3784 | 3856 | 73 | 2 | L |  |  |
| tRNA<sup>Met</sup> (CAT) | 3859 | 3927 | 69 | 0 | H |  |  |
| NADH2 | 3928 | 4971 | 1044 | -2 | H | ATA | TAG |
| tRNA<sup>Trp</sup> (TCA) | 4970 | 5037 | 68 | 5 | H |  |  |
| tRNA<sup>Ala</sup> (TGC) | 5043 | 5111 | 69 | 1 | L |  |  |
| tRNA<sup>Asn</sup> (GTT) | 5113 | 5185 | 73 | 2 | L |  |  |
| OL | 5188 | 5217 | 30 | -1 | H |  |  |
| tRNA<sup>Cys</sup> (GCA) | 5217 | 5282 | 66 | 0 | L |  |  |
| tRNA<sup>Tyr</sup> (GTA) | 5283 | 5349 | 67 | 1 | L |  |  |
| COX1 | 5351 | 6895 | 1545 | -3 | H | ATG | TAA |
| tRNA<sup>Ser</sup> (TGA) | 6893 | 6961 | 69 | 11 | L |  |  |
| tRNA<sup>Asp</sup> (GTC) | 6973 | 7039 | 67 | 0 | H |  |  |
| COX2 | 7040 | 7723 | 684 | 3 | H | ATG | TAA |
| tRNA<sup>Lys</sup> (TTT) | 7727 | 7793 | 67 | 1 | H |  |  |
| ATP8 | 7795 | 7998 | 204 | -43 | H | ATG | TAA |
| ATP6 | 7956 | 8636 | 681 | -1 | H | ATG | TAA |
| COX3 | 8636 | 9420 | 785 | -1 | H | ATG | TAa |
| tRNA<sup>Gly</sup> (TCC) | 9420 | 9488 | 69 | -3 | H |  |  |
| NADH3 | 9486 | 9835 | 350 | 0 | H | ATA | TAa |
| tRNA<sup>Arg</sup> (TCG) | 9836 | 9905 | 70 | 0 | H |  |  |
| NADH4L | 9906 | 10202 | 297 | -7 | H | ATG | TAA |
| NADH4 | 10196 | 11573 | 1378 | 0 | H | ATG | Taa |
| tRNA<sup>His</sup> (GTG) | 11574 | 11642 | 69 | 0 | H |  |  |
| tRNA<sup>Ser</sup> (GCT) | 11643 | 11701 | 59 | 1 | H |  |  |
| tRNA<sup>Leu</sup> (TAG) | 11703 | 11772 | 70 | -9 | H |  |  |
| NADH5 | 11764 | 13593 | 1830 | -17 | H | ATA | TAA |
| NADH6 | 13577 | 14101 | 525 | 3 | L | ATG | TAA |
| tRNA<sup>Glu</sup> (TTC) | 14105 | 14173 | 69 | 4 | L |  |  |
| CYTB | 14178 | 15317 | 1140 | 0 | H | ATG | AGA |
| tRNA<sup>Thr</sup> (TGT) | 15318 | 15386 | 69 | 1 | H |  |  |
| tRNA<sup>Pro</sup> (TGG) | 15388 | 15453 | 66 | 268 | L |  |  |
| CR | 15454 | 16828 | 1388 | / | H |  |  |

**Supplementary Table A4:**
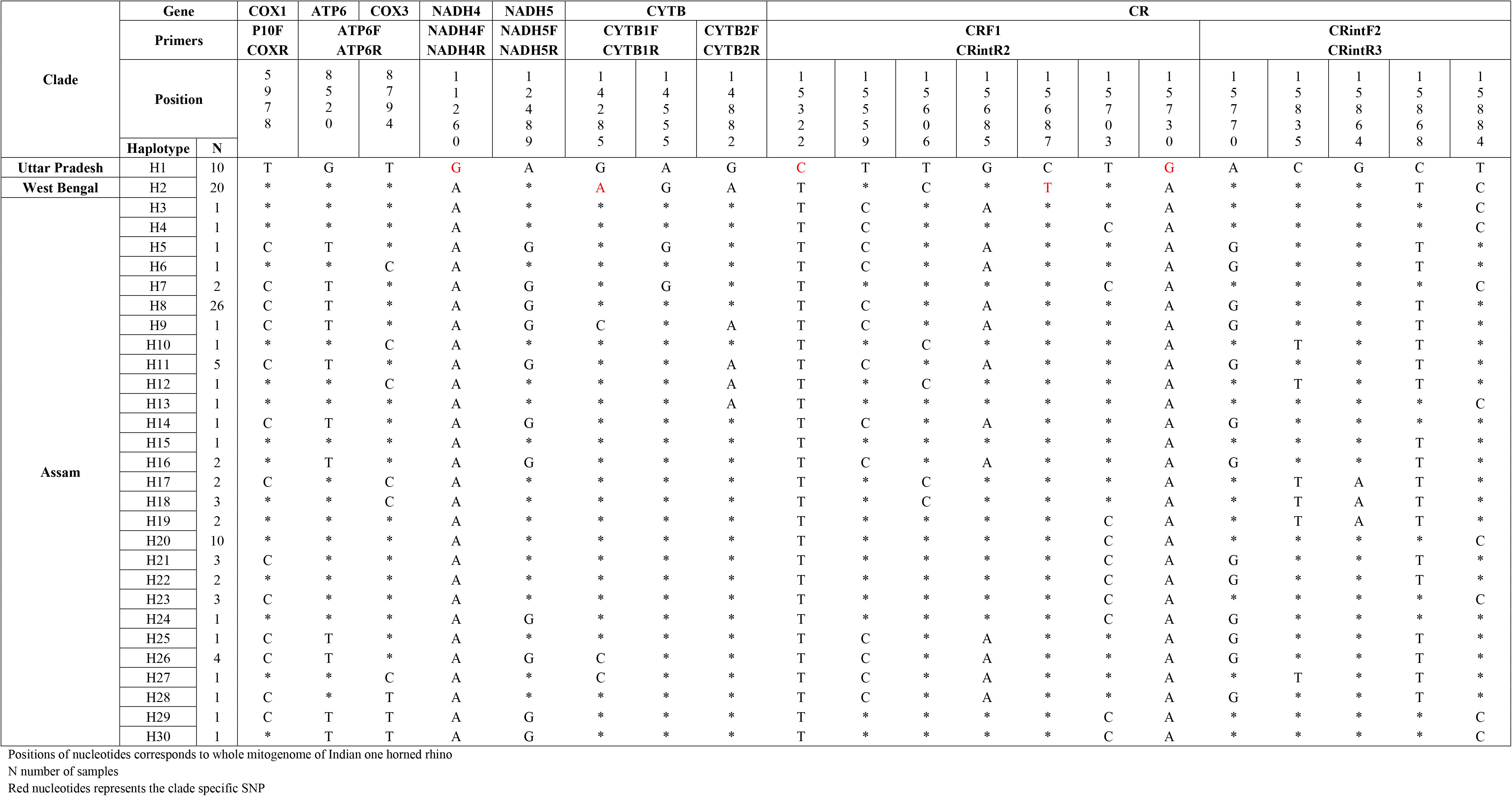
Details of the variable sites based on concatenated sequence of 2531bp of Indian one horned rhino mtDNA haplotypes

